# Analysis of Protein Cysteine Acylation Using a Modified Suspension Trap (Acyl-Trap)

**DOI:** 10.1101/2024.03.23.586403

**Authors:** Michael T. Forrester, Jacob R. Egol, Aleksandra Tata, Purushothama Rao Tata, Matthew W. Foster

**Affiliations:** Division of Pulmonary, Allergy and Critical Care Medicine, Duke University School of Medicine, Durham, NC, 27710, USA; Department of Cell Biology, Duke University School of Medicine, Durham, NC, 27710, USA; Duke Regeneration Center, Duke University School of Medicine, Durham, NC, 27710, USA; Proteomics and Metabolomics Core Facility, Duke University School of Medicine, Durham, NC, 27710, USA

**Keywords:** Suspension trapping, S-Trap, Swisspalm, cysteine proteomics, chemoproteomics *S*-palmitoylation, *S*-acylation, thioester, TMT, isobaric labeling, Fragpipe

## Abstract

Proteins undergo reversible *S*-acylation via a thioester linkage in vivo. *S*-palmitoylation, modification by C16:0 fatty acid, is a common *S*-acylation that mediates critical protein-membrane and protein-protein interactions. The most widely used *S*-acylation assays, including acyl-biotin exchange and acyl resin-assisted capture, utilize blocking of free Cys thiols, hydroxylamine-dependent cleavage of the thioester and subsequent labeling of nascent thiol. These assays generally require >500 micrograms of protein input material per sample and numerous reagent removal and washing steps, making them laborious and ill-suited for high throughput and low input applications. To overcome these limitations, we devised “Acyl-Trap”, a suspension trap-based assay that utilizes a thiol-reactive quartz to enable buffer exchange and hydroxylamine-mediated *S*-acyl enrichment. We show that the method is compatible with protein-level detection of *S*-acylated proteins (e.g. H-Ras) as well as *S*-acyl site identification and quantification using “on trap” isobaric labeling and LC-MS/MS from as little as 20 micrograms of protein input. In mouse brain, Acyl-Trap identified 279 reported sites of *S*-acylation and 1298 previously unreported putative sites. Also described are conditions for long-term hydroxylamine storage, which streamlines the assay. More generally, Acyl-Trap serves as a proof-of-concept for PTM-tailored suspension traps suitable for both traditional protein detection and chemoproteomic workflows.

**GRAPHICAL ABSTRACT:** 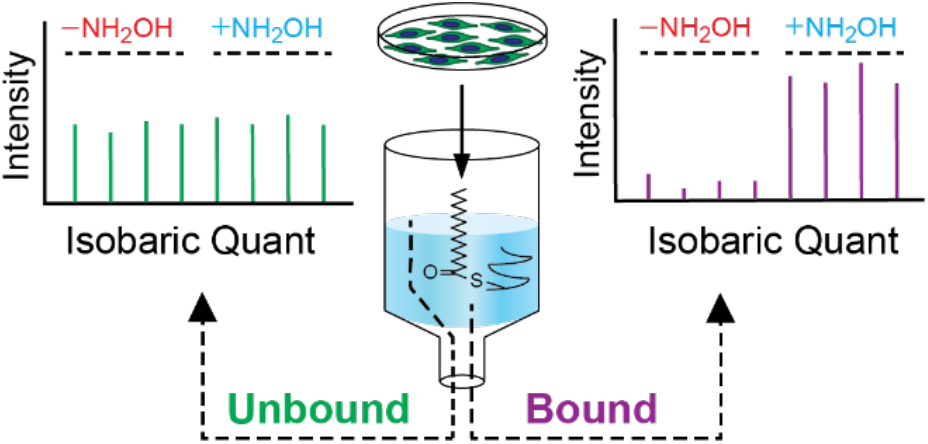

## INTRODUCTION

Protein fatty acylation is a ubiquitous post-translational modification that involves esterification of a Cys thiol with a saturated lipid typically 16C in length (*S*-palmitoylation), imparting significant hydrophobicity onto the modified protein. Starting with the discovery of viral *S*-palmitoylation in 1979 by radiolabeling and autoradiography^1^, decades of methodological advancement have established roles of *S*-palmitoylation in a panoply of biological processes^2-4^. Examples include membrane trafficking of proto-oncogene Ras^5,6^ and Src^7,8^ isoforms, β-adrenergic receptor signaling^9-11^, vesicle/vacuole fusion^12,13^ and proper assembly of respiratory viruses^14-16^. Large scale computational analyses suggest that aberrant *S*-palmitoylation may play a role in cancer and neurologic disorders^17^ and pharmacological studies targeting the *S*-palmitoylation machinery show promise for the future development of antiviral^18,19^ and antineoplastic^20,21^ therapies.

Introduced in 2004, Acyl-biotin exchange (ABE), which utilizes blocking of free thiols, NH_2_OH-based cleavage of the Cys thioester and subsequent reaction with biotin disulfide for biotin-avidin enrichment, was a major advance over the use of radiolabeled palmitate^22,23^. Acyl-RAC, which we described in 2011, replaced Cys-biotinylation and avidin enrichment with a single “resin-assisted capture” step using a Sepharose-immobilized reactive disulfide, thus enabling facile *S*-acyl site identification and quantification^24^. In conjunction with lipid labeling approaches^25-28^, ABE and Acyl-RAC have become mainstay assays for targeted *S*-palmitoylation studies and *S*-acyl proteomics^29^.

While these methodological advances have been paramount for driving the *S*-acylation field, ABE and Acyl-RAC rely on laborious protein precipitation steps for reagent removal and usually require extensive washing of streptavidin or thiopropyl sepharose. These assays are best accomplished with fairly large quantities of protein per sample (typically > 500 μg) and can require several days to complete. Multiple precipitation steps of ABE are time-consuming and can result in selective protein loss^30,31^ that can be partly mitigated by resin-assisted capture or chemical scavenging of thiol blocking reagent^32^. Methods that require less input material and require fewer steps than current gold standards could further accelerate drug discovery efforts and make *S*-acylation a more routine analysis for multi-omic and systems biology investigations.

Since its introduction Zougman et al. in 2014^33^, suspension trapping has gained popularity for ease of detergent removal and efficient proteolysis, thus supporting many bottom-up proteomic workflows^34,35^. Similar to ABE and Acyl-RAC, suspension trapping utilizes protein denaturation and cysteine alkylation followed by protein precipitation as the initial steps. However, rather than using centrifugation to recover the precipitated protein, suspension trapping utilizes a quartz plug to physically capture protein and facilitate exhaustive washing, thus enabling the processing of very small amounts of protein. Given a report demonstrating cysteinyl-peptide enrichment on a chemically modified quartz trap^36^ and our experience with Acyl-RAC, we conceived a *S*-acyl assay utilizing thiol-reactive quartz in lieu of thiol-reactive resin or biotinylating reagent. Such an approach could combine the many advantages of suspension trapping with established *S*-acyl enrichment chemistry while serving as a proof-of-concept for other chemoproteomic workflows using derivatized suspension traps.

## EXPERIMENTAL PROCEDURES

Materials – All chemicals were obtained from Sigma-Aldrich unless otherwise specified. Acid labile surfactant (ALS-1)^37^ was synthesized by the Duke Small Molecule Synthesis Facility. Sequencing-grade modified trypsin was from Promega.

### Synthesis of pyridyl disulfide quartz (PDQ)

Up to eight 4.7 cm diameter MK360 filters (Ahlstrom) were placed into a 60 ml “TraceClean” glass jar (Avantor). Ten milliliters of xylenes was added followed by 2 ml of (3-Mercaptopropyl)trimethoxysilane (MPTS) to achieve final concentration of ∼1 M MPTS. After 12-24 h incubation on a rotating platform at room temp, the filters were transferred to a Büchner funnel and washed five times with at 200 ml of xylenes. The washed filters were transferred to a new jar containing 200 mg 2-pyridyl disulfide (final PDS 100 mM) in 10 ml xylenes. After 1 h at room temp, the filters were washed five times with 50 ml xylenes and 10 times with 50 ml isopropanol, after which the filters change from semi-translucent appearance to opaque white. The resulting PDQ filters were placed onto aluminum foil in a single layer and dried at 50 °C (on heat block or in an oven) for 30-60 min. The PDQ filters were stored in a 6 cm polystyrene cell culture dish at room temp until use.

### Construction of PDQ traps

Three plugs of PDQ were captured into a clipped P200 pipette tip (∼1/16th inch i.d.) and gently packed in a clean P200 tip using 1/16th inch o.d. PEEK tubing (Figure S1). This process was performed in triplicate for each P200 tip, for a final of 9 PDQ “plugs” per trap (Figure S2). Traps were stored at room temp.

### Cell culture and murine tissue harvesting

MLE12 cells were obtained from the Duke Cell Culture Facility and cultured in HITES media (50:50 Dulbecco’s medium/Ham’s F12 supplemented with 0.005 mg/ml insulin, 0.01 mg/ml transferrin, 30 nM sodium selenite, 10 nM hydrocortisone, 10 nM β-estradiol, 2 mM GlutaMAX and 2% FBS). Cells were grown to 80% confluency in 6-well dishes or 10 cm plates and washed twice with cold PBS prior to lysis. Mice (C57BL/6) were euthanized by carbon dioxide followed by bilateral thoracotomy and perfused with 5-10 ml of PBS through the right ventricle. Cerebri were harvested, snap frozen and stored at -80 °C. All mouse handling was in accordance with the Duke University Institutional Animal Care and Use Committee (IACUC) and NIH guidelines.

### Acyl-Trap protocol for Western blotting

All buffers were between pH 7.2 – 7.4, and centrifugation was at 3000 *xg* for 1 min unless otherwise indicated. Following collection in PBS, HEK293 cells were lysed in 50 mM HEPES, 5 mM EDTA, 0.5% Triton X-100 via probe sonication and clarified by centrifugation at 1000 *xg* for 5 min. Sample concentrations were measured by detergent-compatible Bradford assay (Sigma). Fifty micrograms were brought to 40 μL with 50 mM HEPES, 5 mM EDTA then 10 μL of 20% SDS is added to achieve a final of 2% SDS (w/v). Following addition of 1 μL fresh 1M NEM in MeOH (final 20 mM NEM), samples were incubated at 50 °C for 1 h with frequent vortexing. Three volumes (150 μL) of room temperature MeOH were added followed by vortexing and incubation at room temp for 10 min. The slurry was loaded to a PDQ trap and centrifuged until all liquid has spun through the trap. The traps were washed four times with 200 μL MeOH. To the traps were added 15 μL of 100 mM HEPES, 5 mM EDTA and 1% SDS, with or without NH_2_OH (0 for control, up to 500 mM for the +NH_2_OH sample). Traps were incubated at room temp for 30 min and centrifuged to recover the reaction solution. This process was repeated with another 15 μL of reaction solution and 30 min room temp incubation. Traps were washed four times with 200 μL of 50 mM HEPES, 5 mM EDTA, 1% SDS. For on-PDQ biotinylation, 40 μL of PBS pH 8.0 containing 5 mM Biotin-NHS was added to each trap. After 30 min, the trap was spun and another biotinylation reaction performed for 30 min followed by a wash with 200 μL of PBS x2. For elution, traps are moved into a clean microfuge tube and incubated with 20 μL of 50 mM HEPES, 5 mM EDTA, 1% SDS containing 20 mM DTT. After 20 min, eluates were collected by centrifugation. This step was repeated in the same microfuge tube for total 40 μL eluate (bound material), which was then subjected to SDS-PAGE.

### Acyl-Trap protocol for LC-MS

Cells or tissues were lysed in 50 mM HEPES, 5 mM EDTA and 0.5% Triton X-100 containing cOmplete Mini protease inhibitor (Roche) via 3 rounds of probe ultrasonication. Crude lysates were centrifuged at *2000 g* for 5 min to remove intact cells and large fragments (e.g., extracellular components, nuclear chromatin). Detergent-compatible Bradford (Thermo) was performed on the remaining lysate. Twenty to 50 μg of lysate protein was brought to 50 μL volume with lysis buffer including final 2% SDS and 20 mM NEM (from a 1 M stock in EtOH). After heating at 50 °C for 30 min on a Thermomixer (Eppendorf) at 1000 rpm, 3 volumes of MeOH were added to each sample, incubated for 10 min at room temp and loaded onto a PDQ trap via centrifugation. Traps were washed four times with 200 μL MeOH. To each trap was added 20 μL of digestion solution (50 mM HEPES, 5 mM EDTA, 0.5% ALS-I and 2 μg of trypsin) containing either 200 mM NaCl or 200 mM NH_2_OH. Traps were covered with parafilm that was perforated once with a 21G needle, placed into a clean de-capped 1.7 ml microfuge tube and incubated at 42 °C for 1 h using a Thermomixer with heated lid at 0 rpm. Following this proteolysis and capture step, “unbound” material was collected by adding 50 μL of 50 mM HEPES and 5 mM EDTA followed by centrifugation for 2 total collections.

Unbound material was adjusted to 1% TFA and heated at 50 °C for 1 h to decompose ALS-1 followed by centrifugation. Unbound material was subjected to C18 STAGE-Tip based TMT labeling by the method of Myers et al^38^. In brief, five micrograms of peptide digest (based on starting input protein material) was adjusted to 50 μL with 0.2% (v/v) formic acid. Stage tips were equilibrated with MeOH followed by 50% acetonitrile / 0.2% formic acid. Unbound protein was loaded to the stage tip and washed twice with 40 μL 0.2% formic acid. For TMT labelling, TMTpro 16 plex reagents (0.5 mg each, Thermo) were reconstituted in 20 μL DMSO. Two microliters were mixed with 50 μL 100 mM TEAB and immediately added to the C18 stage tip. The tips were centrifuged at 200 *xg* for 5 min, 1000 *xg* for 1 min, then washed twice with 40 μL 0.2% formic acid. Peptides were eluted with 50 μL of 50% acetonitrile/0.2% formic acid, then 50 μL 20 mM ammonium formate in 50% acetonitrile. Each elution was performed via centrifugation at 600 *xg* for 2 min, then 1500 *xg* for 2 min. Eluants were combined and lyophilized.

For processing of the PDQ-bound peptides, proteolyzed traps from above were washed with 200 μL of 50 mM HEPES + 1% (w/v) SDS four times followed by 200 μL 50 mM TEAB twice. For TMT labelling, 4 μL of reconstituted TMTpro reagent was mixed with 16 μL of 50 mM TEAB, immediately added to each PDQ trap, covered with parafilm and incubated at 40 °C for 1 h. Each trap was washed four times with 200 μL of 50 mM TEAB by centrifugation and then eluted twice by incubation with 40 μL TEAB containing 20 mM DTT at room temp for 20 min each. Eluates were recovered by centrifugation, combined, and alkylated with iodoacetamide (2.5 molar excess over DTT) for 30 min at room temp, acidified with 1% final (v/v) TFA and lyophilized.

### Liquid chromatography coupled to tandem mass spectrometry (LC-MS/MS)

PDQ-enriched peptides were reconstituted in 15 μL of 97/2/1 (v/v/v/) H2O/MeCN/TFA, and 4.5 μL was analyzed in duplicate by LC-MS/MS. Unbound fractions were reconstituted in the same buffer at ∼1 μg/μL, and 1.5-2.5 μL was analyzed. LC-MS/MS used a Waters M-Class interfaced to a Thermo Fusion Lumos via a NanoSpray Flex source and CoAnn emitter 20 μm ID, 10 μm Tip ID, 6.35 cm length)). Peptides were trapped on a Symmetry C18 180 μm x 20 mm trapping column (Waters) at 5 μL/min with 99.9/0.1 v/v H_2_O/MeCN and separated on a 75 μm x 25 cm HSS-T3 analytical column (Waters) at 400 nl/min and 55 °C and a gradient of 5-30% MeCN over 90 min. MS acquisition used a 3 s vendor-supplied synchronous precursor selection (TMTPro-SPS-MS3) template method. Briefly, Precursor scans used 120,000 resolution, standard AGC target and auto maximum injection time (IT). Precursors (scan range 400-1600 m/z, charge state 2-6, intensity threshold 5E3) were selected for collision-induced dissociation using 0.7 m/z, “Turbo” ion trap scan with 25 ms max IT. Finally 10 SPS precursors were subjected to higher-energy collision-induced dissociation with 0.7 m/z isolation, normalized collision energy of 55%, 50,000 resolution, 200% AGC and 200 ms max IT. Analysis of unbound fractions also used a FAIMS-Pro interface with a 3 CV method (−40 CV for 1.2 s, -55 CV for 1.2 s, -70 CV for 0.8 s).

### Database searching and isobaric quantification

Raw data were converted to mzml using MS Convert v. 3.0.21193-ccb3e0136 (Proteowizard)^39^ and searched with Fragpipe (v21.1)^40,41^ with a TMT16-MS3 workflow which included the following settings: precursor mass tolerance 20 ppm, fragment mass tolerate 0.8 Da, cleavage rules enzymatic with N-terminal clipping enabled using stricttrypsin and allowing for 2 missed cleavages, peptide mass range 7 – 40 aa and 500 – 5000 Da. A Mus musculus database was downloaded from UniProt (ProteomeID UP000000589) on 1/26/2024 using FragPipe and appended with common contaminants and reverse decoys (total number of entries = 34,560). For bound material, fixed modifications were Lys TMT (+304.20715 Da) and Cys carbamidomethylation (+57.02146 Da); variable modifications were protein N-term acetylation (+42.0106 Da), protein N-term myristoylation (+210.19836 Da), peptide N-term TMT (+304.20715 Da) and Ser TMT (+304.20715 Da) allowing for maximum 2 variable mods per peptide. For isobaric settings, label type = TMT-16, quant level = 3, mass tolerance = 20 ppm, group by = peptide, normalization = none, PTM mod tag = C(57.02146), min site probability 0. For unbound material, fixed modifications were Lys TMT (+304.20715 Da) and Cys NEM adduct (+125.047676 Da); variable modifications were N-term acetylation (+42.0106 Da), N-term TMT (+304.20715 Da) and Ser TMT (+304.20715 Da) allowing for maximum 5 variable mods per peptide. For isobaric settings, label type = TMT-16, quant level = 3, mass tolerance = 20 ppm, group by = protein, normalization = MD, PTM mod tag = none, min site probability -1.

### Data post-processing and analysis

Fragpipe TMT output data were read into Rstudio (Version 2023.09.0+463). For unbound data, missing values in the protein-level TMT output were imputed with the lowest 5% value. For bound samples, “multisite” TMT data were subjected to quantile normalization with grouping (n=4 each minus and plus NH_2_OH) followed by imputation with the lowest 3% value on the no-NH_2_OH replicates if 2 or more values were missing. In other words, if 3 of 4 no-NH_2_OH replicates had values, the 4^th^ missing value was not imputed. For bound sites with corresponding protein quantification data in the unbound, the bound data were subjected to log2 subtraction of the unbound intensity from the corresponding replicate (to account for any relative changes in protein abundance). Log2 fold change (-/+ NH_2_OH) was calculated by subtracting the mean log2 intensity of minus NH_2_OH replicates from the mean log2 intensity of plus NH_2_OH replicates. For peptides containing at least 3 values for each group of replicates, a paired t-test was performed followed by FDR thresholding using the Benjamini-Hochberg method.

### Database matching and GO analysis

UniProt mouse proteome (Release 2023_03) and Swisspalm (swisspalm.org, version 4, 2022-09-03) were imported into Rstudio. Sites of *S*-palmitoylation in UniProt were identified in column “Lipidation” and extracted with the stringr package to identify 462 unique sites. Swisspalm mouse data (total 4381 sites of *S*-palmitoylation) were filtered for column “large_scale = No” to identify 310 sites that were the results of small-scale targeted studies. By combining UniProt and confirmed Swisspalm data, 601 unique *S*-palmitoylated Cys sites were employed as the experimentally confirmed “small-scale” dataset analogous to PhosphoSite Low Throughput (LTP) classification^42^. All remaining Swisspalm data were designated as High Throughput (HTP) in a manner analogous to the PhosphoSite annotation. Matching between the experimental data and these databases was achieved with the stringr package.

Over-representation analysis (ORA) was performed with the enrichGO function within the clusterProfiler package^43^ by comparing genes lists for the proteins containing enriched high probability Cys sites to a universal gene list containing the above proteins along with unenriched proteins and proteins identified in the unbound sample that were not represented in the enriched (in other words, were present in the lysate but not detected in the bound). Within each list, duplicate Cys sites were represented by only 1 unique gene. Settings for the ORA included: OrgDb = org.Mm.eg.db, ont = “ALL”, pAdjustMethod = “BH”, pvalueCutoff = 0.05, qvalueCutoff = 0.05).

## RESULTS AND DISCUSSION

### Conception and Implementation of Acyl-Trap Workflow

Protein *S*-acyl detection frequently employs a 3-step strategy: 1) free thiol alkylation (i.e., blocking), 2) NH_2_OH-mediated thioester cleavage and 3) capture of newly liberated thiol with either an affinity reagent (e.g. biotin-HPDP) or thiol-reactive Sepharose in the case of Acyl-RAC. To develop Acyl-Trap, we envisioned an analogous strategy that replaces thiol-reactive biotin or Sepharose with a thiol-reactive suspension trap (pyridyl-disulfide quartz, PDQ). Illustrated in Figure 1, Acyl-Trap commences with protein denaturation and blocking with N-ethylmaleimide (NEM), followed by MeOH-based precipitation and trapping. Trapped proteins are washed to remove excess NEM and SDS, then resolubilized in the presence or absence of NH_2_OH. In the Western blot protocol, proteins are covalently captured onto PDQ via NH_2_OH-liberated Cys thiols and eluted with reductant. In the LC-MS protocol, proteins are subjected to simultaneous proteolysis and peptide-level capture. Akin to a traditional suspension trap, unbound or “flow through” material is optionally collected for quantification of protein abundance. The PDQ-bound peptides are subjected to isobaric (TMT) labelling, eluted with reductant, combined and alkylated.

**Figure 1.**
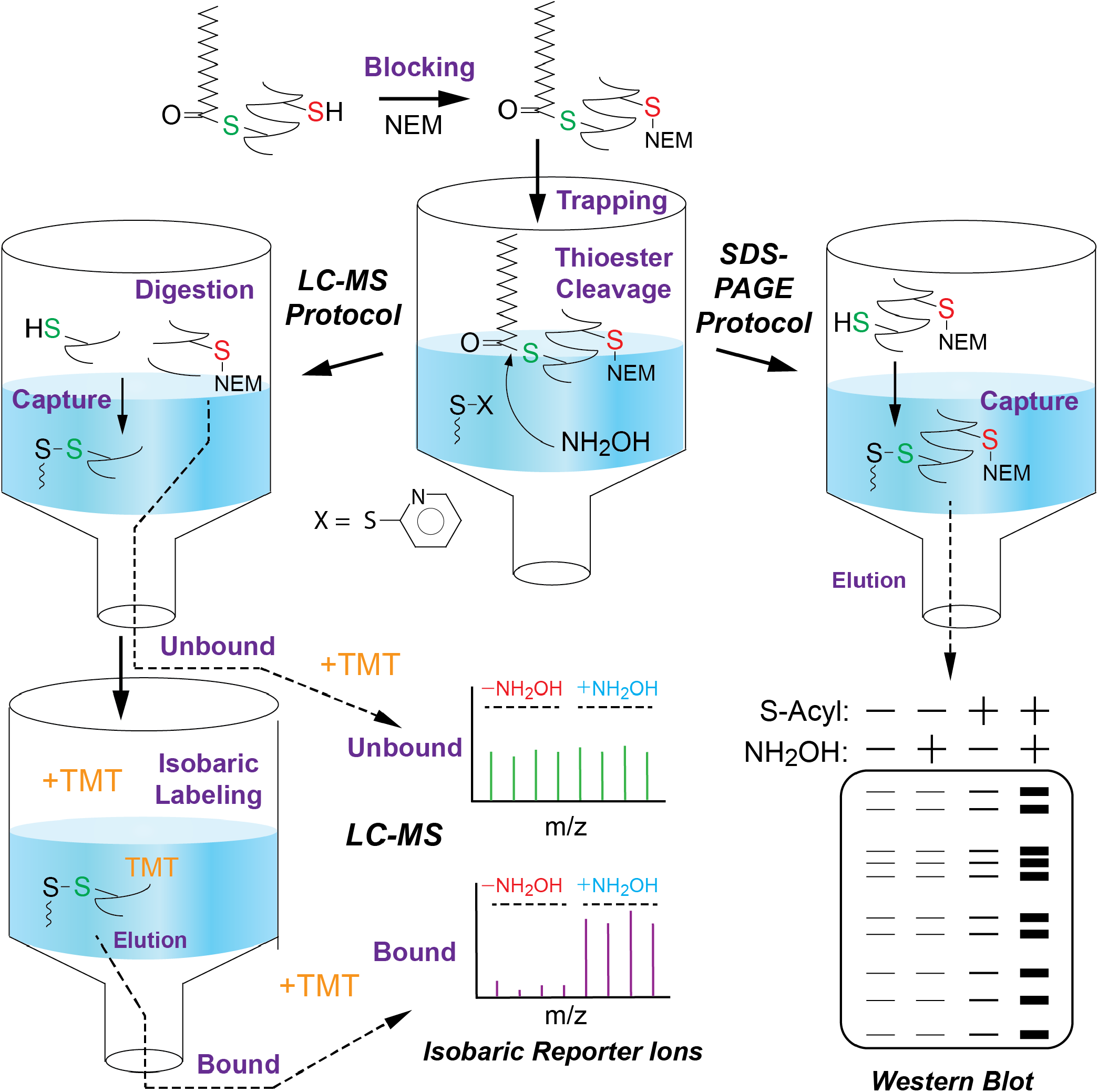
Schematic overview of Acyl-Trap. Protein-containing lysates are first alkylated (“Blocking”) with *N*-ethylmaleimide (NEM) in the presence of SDS. Following addition of MeOH, the insolubilized protein slurry is physically trapped onto the pyridyl disulfide quartz (PDQ), and NEM/SDS are removed by centrifugation and MeOH washing. A solution of SDS (Western Blot Protocol), or trypsin and acid-labile surfactant (LC-MS Protocol), are added to the trapped proteins along with hydroxylamine (NH_2_OH), which cleaves thioesters to free thiols, resulting in covalent “capture” of proteins/peptides by the electrophilic 2-thiopyridyl disulfide of PDQ. Unbound proteins are removed by washing and bound proteins eluted with reductant. For MS, unbound peptides are collected and subjected to cleanup (i.e. NH_2_OH removal) and TMT labeling on C18 StageTips before pooling and analysis by LC-MS/MS. PDQ-bound peptides are subjected to TMT labeling, elution with reductant, pooling, alkylation and LC-MS/MS analysis.

### Synthesis of Pyridyl Disulfide Quartz (PDQ) and Assembly of Thiol-Reactive Suspension Traps

In order to develop Acyl-Trap, we first focused on possible synthetic routes to a thiol-reactive quartz. The synthesis described by Zougman et al.^36^ utilized the amine-thiol crosslinking agent succinimidyl 3-(2-pyridyldithio)propionate (SPDP), which is expensive and limits the large scale synthesis of PDQ. In addition, MK360 filters appeared to delaminate in acetone, the reported reaction solvent. We developed a simple alternative synthetic scheme, performing silanization with a mercaptosilane followed by oxidation with 2-pyridyl disulfide (Figure S1A). The silanization is performed similarly to published silica-based reactions^44,45^ in organic media (toluene or xylenes) using glass chambers and a Buchner funnel (Figure S1B). To create a trap, small discs of dried PDQ are packed into a P200 pipette tip (Figures S1C, S2A-C). Like other tip-based protocols, PDQ traps are amenable to processing/centrifugation in small or large batches (Figures S1D, S2D-E). The mercaptosilane-derivatized quartz shows reactivity with 5,5′-dithiobis-2-nitrobenzoic acid (DTNB) generating an intensely yellow color, which is expectedly lost upon conversion to PDQ (Figure S1E). The PDQ is stored dry at room temp until use.

### Unique Methodological Considerations of Acyl-Trap

In developing Acyl-Trap, we made several modifications to the typical suspension trapping protocol requiring some validation or explanation. Thioesters are particularly sensitive to acid- or base-catalyzed hydrolysis^46^. However, the typical S-Trap protocol uses phosphoric acid to presumably facilitate SDS neutralization. To avoid conditions that could compromise thioester stability, we evaluated the efficiency of protein trapping under neutral pH conditions. Shown in Figure S3A, 3-5 volumes of MeOH or EtOH provided efficient and consistent protein trapping from both SDS and GdnCl-denatured samples. Thus, for the purposes of Acyl-Trap it is acceptable to maintain neutral pH during trapping.

During Acyl-RAC or ABE, blocking and NH_2_OH-mediated capture are performed at the protein-level followed by subsequent proteolysis for the purposes of identifying putative *S*-acylated sites. In contrast, for downstream LC-MS/MS, Acyl-Trap combines proteolysis, NH_2_OH-mediated thioester cleavage and thiol capture into a single step. To ensure that NH_2_OH does not impact trypsin activity and therefore affect the degree of peptide recovery, the colorimetric BAPNA assay was performed in the presence of NH_2_OH (Figure S3B). No appreciable inhibition or augmentation of BAPNA hydrolysis by NH_2_OH was measured, demonstrating that NH_2_OH-based thioester cleavage and trypsin activity are chemically compatible.

### Analysis of H-Ras and Global Protein *S*-Acylation by Western Blot-Compatible Acyl-Trap

To assess the ability of Acyl-Trap to process intact *S*-acyl proteins for analysis by SDS-PAGE, 40 μg of HEK293 lysate was subjected to Acyl-Trap coupled with on-PDQ biotinylation. Eluants were visualized with avidin-HRP. In a manner similar to ABE and Acyl-RAC, Acyl-Trap revealed a marked NH_2_OH-dependent augmentation in covalent protein capture (Figure 2A). These results also establish that intact proteins are capable of undergoing elution from PDQ in the presence of SDS and DTT, and thus do not require proteolysis for their recovery.

**Figure 2.**
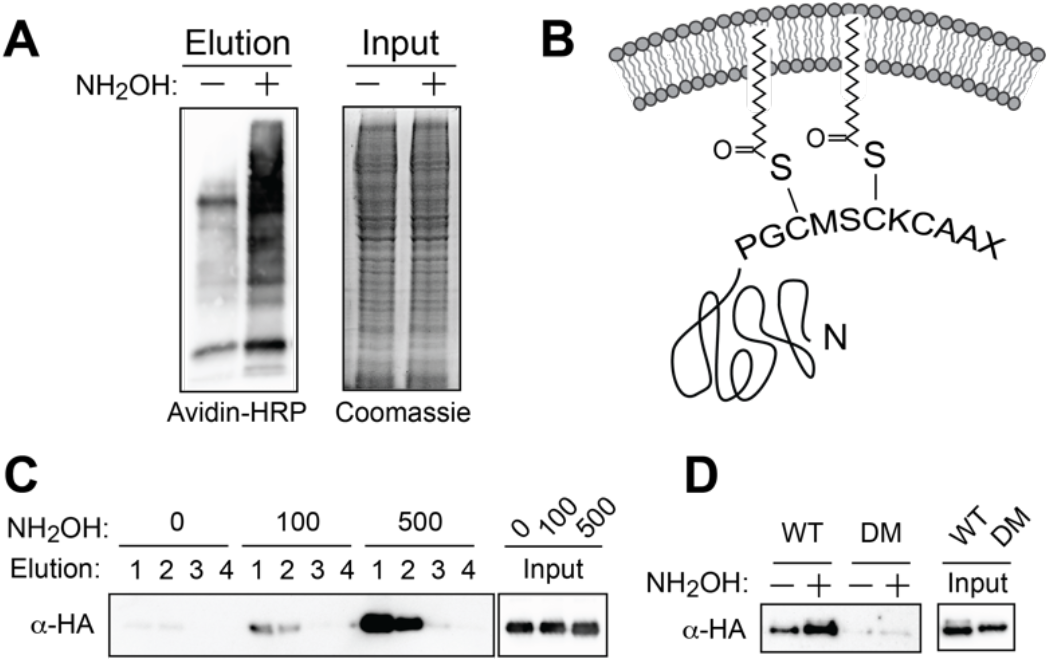
Acyl-Trap in conjunction with SDS-PAGE / Western Blot analysis. (A) HEK293 cells (40 μg per sample) were subjected to Acyl-Trap, on-quartz biotinylation and chemiluminescent detection with Avidin-HRP. Prior to trap loading, 10 micrograms of each blocked sample were collected and analyzed by Coomassie staining. (B) Schematic of H-Ras C-terminal palmitoyl Cys sites. The final 4 amino acids (CAAX motif) undergo Cys prenylation in the form of a stable thioether, proteolysis and carboxymethylation. (C) HEK293 cells were transfected with HA-tagged H-Ras and 40 μg of sample were subjected to Acyl-Trap. Prior to trap loading, 10 μg of each blocked sample were collected for input. Detection of *S*-acylated H-Ras was assessed by performing sequential 20 μl elutions with DTT followed by western blotting with anti-HA antibody. (D) Comparison of wild type (WT) and non *S*-palmitoylated C181S/C184S double mutant (DM) H-Ras. Results are representative of at least n=4 separate experiments.

Proto-oncogene H-Ras is a well-established target of *S*-palmitoylation on two C-terminal Cys sites^6^ (Figure 2B) and was employed for the development of Acyl-RAC. To examine the feasibility of Acyl-Trap to enrich this model *S*-palmitoylated protein, HEK293 cells were transfected with HA-tagged H-Ras and 40 μg of lysate subjected to Acyl-Trap. As shown in Figure 2C, enrichment of H-Ras was markedly augmented by NH_2_OH and two rounds of reductive elution (20 μl each) were sufficient for H-Ras recovery. Enrichment was dependent on *S*-palmitoylation, as assessed by comparing wild type vs C181S/C184S double mutant (DM) H-Ras (Figure 2D). Distinct from ABE and Acyl-RAC, Acyl-Trap circumvents labor intense protein resuspension steps and allows processing of much smaller quantities of material. To our knowledge this is the first demonstration of suspension trapping for the purpose of covalent PTM enrichment at the level of whole protein (i.e. in the absence of proteolysis).

### Application of Acyl-Trap to Mouse Brain and Lung Epithelial Cell Line

To test the generalizability of Acyl-Trap, mouse cerebrum (50 μg per sample) and murine lung epithelial cell line MLE12 (20 μg per sample) were subjected to Acyl-Trap as separate 8-plex TMT experiments utilizing 4 channels each for the non-NH_2_OH and plus-NH_2_OH replicates. An evaluation of precursor ions for the presence of a Cys residue revealed enrichment of Cys-containing peptides with 75-90% of peptides in the bound material having at least 1 Cys residue compared to 7-11% in the unbound material (Figure S4). These findings are on par with global Cys peptide enrichment reported by Zougman et al^36^.

Amine reactive isobaric labels (e.g., TMT) rely on a nucleophilic peptide amine reacting with an *N*-hydroxy succinimide (NHS) ester to form a stable amide bond. Analysis of precursor ion data – subjected to a search with variable N-term and Lys-TMT modification – revealed a modest underlabeling of PDQ-immobilized peptides, exemplified by non-Lys containing (i.e. C-terminal Arg) peptides where 13-16% of these did not undergo TMT modification. In contrast, C18-immobilized TMT labeling^38^ exhibited only 2-5% under-labeling on similar non-Lys containing peptides (Figure S5A) suggesting a slight decrease in amine reactivity for PDQ-immobilized peptides. Conversely, over-labelling—as defined by TMT modification to Ser/Thr/Tyr/His side chains^47^ —was slightly lower with on-quartz TMT labeling versus on-C18 (Figure S5B). Collectively, these findings demonstrate that Acyl-Trap enriches for Cys-containing peptides and is compatible with on-quartz TMT labeling, with under/overlabeling within acceptable ranges.

Analysis of the TMT quantitative data from both Brain and MLE12 samples revealed a wide range of NH_2_OH-dependent enrichment (Figure 3A-B). Peptides were assigned to enriched high probability (Log2FC > 0.59 and adjusted p-value < 0.05), enriched low probability (Log2FC > 0.59 and adjusted p-value > 0.05) and unenriched (red, Log2FC < 0.59). Utilizing these cutoffs, the brain experiment assigned 1110 and 183 peptides as high and low probability enriched, respectively, while 717 peptides were considered unenriched. In the MLE12 experiment, 948 and 242 peptides were assigned as high and low probability, respectively, while 624 peptides were considered unenriched.

**Figure 3.**
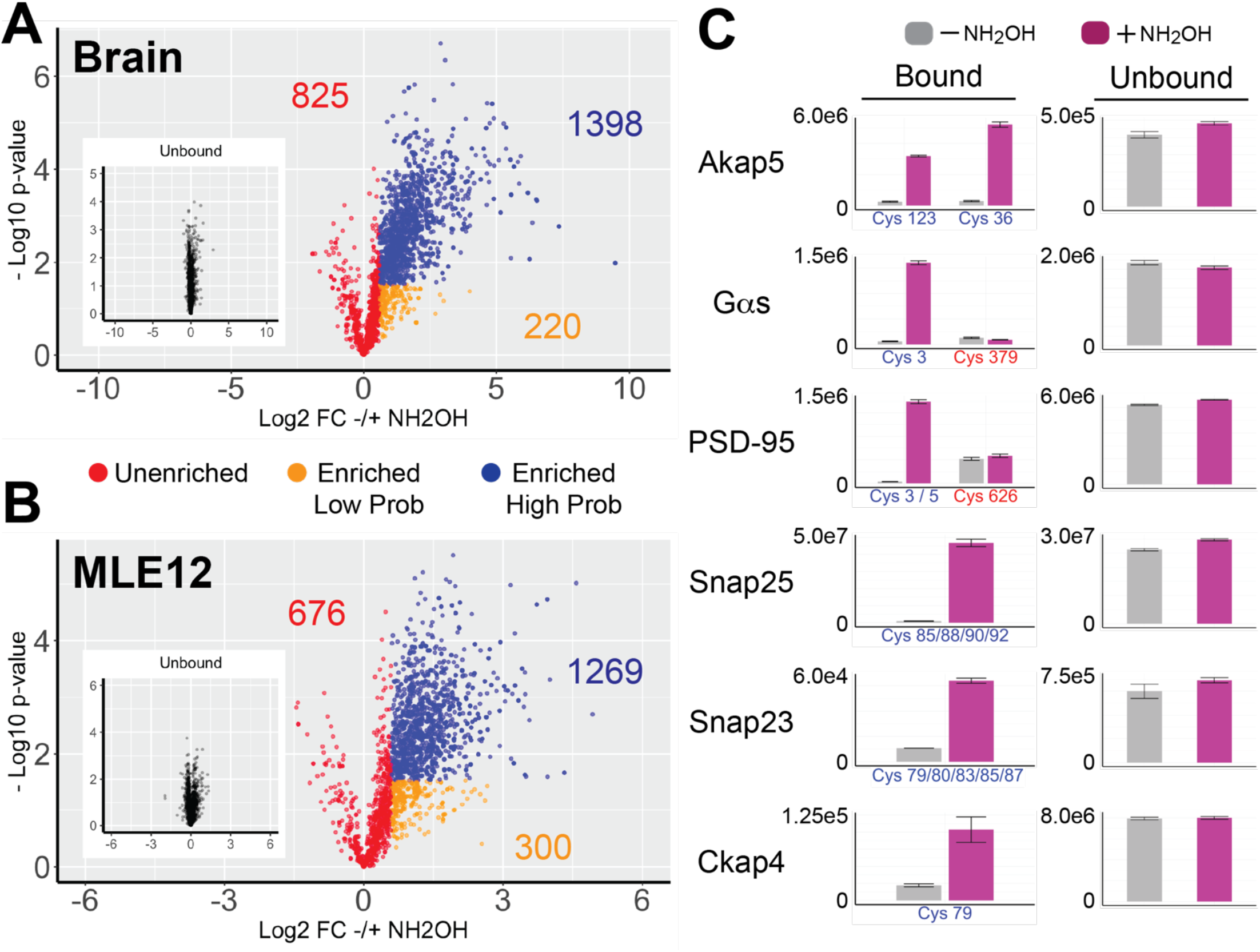
Acyl-Trap coupled with isobaric (TMT) labeling and LC-MS/MS. Volcano plot of Acyl-Trap data from (A) mouse brain using 50 micrograms per replicate and (B) mouse lung epithelial cell line MLE-12 using 20 micrograms per replicate. Each data point represents a unique peptide along with the number of unique Cys sites in corresponding color. Results from the bound material (i.e., subjected *S*-acyl enrichment) are colored according to 3 assignments: enriched high probability (blue, Log2FC > 0.59 and adjusted p-value < 0.05), enriched low probability (orange, Log2FC > 0.59 and adjusted p-value > 0.05) and unenriched (red, Log2FC < 0.59). Shown in the inset are the unbound data with identical x- and y-axes for comparison. (C) Comparison of NH_2_OH-dependence (left) for well-established *S*-acyl sites from brain (Akap5, heterotrimeric G-protein Gαs, PSD-95, and Snap-25) and MLE-12 cells (Snap23 and Ckap4). Y-axes are relative intensities for raw (i.e. non-log2 transformed) data. Data from minus NH_2_OH and plus NH_2_OH replicates are colored in gray and magenta, respectively. Sites known to undergo *S*-palmitoylation have a blue x-axis label, whereas identified sites not known to be *S*-palmitoylated are labeled in red. Corresponding protein abundances from the unbound fraction are shown in the right column.

The larger proportion of enriched low probability in MLE12 compared to brain likely reflects the slightly larger median coefficient of variation in MLE12 compared to brain data (Figure S6), along with less complete TMT quantitation as 87.3% of the MLE12 multi-site data were represented in 8 out of 8 TMT channels whereas for brain this was 97.1%. These differences are likely attributable, at least in part, to the significantly less MLE12 input used for each replicate (20 vs 50 μg for brain). In both experiments, no more than 1% of precursor ions contained an oxidized Met side chain (Figure S7) suggesting that the Acyl-Trap protocol does not lead to appreciable Met oxidation and allowing for this variable modification to be omitted from database searching.

Focusing on established *S*-acylated proteins revealed the ability of Acyl-Trap to identify well-validated *S*-acyl sites (Figure 3C). For example, synaptosomal-associated protein 25 (SNAP-25) and orthologue SNAP-23 – important mediators of plasma membrane fusion – rely on a cluster of 4 or 5 Cys residues, respectively, for proper trafficking and membrane association^13,48,49^. Accordingly, in brain and MLE-12 cells, the corresponding Cys-containing peptides from SNAP-25 and SNAP-23 were enriched in a NH_2_OH-dependent manner, respectively. Other examples include A-kinase anchor protein 5 (Akap5), heterotrimeric GTPase Gαs and post-synaptic density protein 95 (PSD-95, DLG4). In the cases of Gαs^50^ and PSD-95^51^, *S*-acylation is known to be restricted to conserved N-terminal Cys residues. Consistent with these well-established experiments, Acyl-Trap exhibits NH_2_OH-dependent enrichment towards the expected N-terminal Cys-containing peptides, yet two other identified Cys-containing peptides not known to undergo *S*-acylation were unchanged by the presence of NH_2_OH.

### Matching of Acyl-Trap Data to Palmitoylation Databases and *S*-Acyl Site Discovery

For the purposes of evaluating Acyl-Trap, a list of 601 curated / validated palmitoyl Cys sites was assembled by combining *S*-palmitoyl data from UniProt with small scale data in SwissPalm^52^. This *S*-palmitoyl low throughput (LTP) site is analogous to PhosphoSite designation of low vs. high throughput data^42^. As shown in Figure 4A, results from Acyl-Trap showed a total of 42 and 15 matches to these sites between Brain and MLE-12 experiments, respectively. Within these matches, Acyl-Trap assigned 41/42 (98%) from brain and 15/15 (100%) from MLE12 as enriched by NH_2_OH. In the brain and MLE-12 experiments, these site matches exhibited a mean Log2FC ± NH_2_OH of 3.90 and 2.87, along with mean adjusted p-values of 0.019 and 0.017, respectively. Collectively these findings demonstrate that, when Acyl-Trap identifies peptides bearing well-validated sites of *S*-palmitoylation, the method correctly assigns them to the high-probability enriched group.

**Figure 4.**
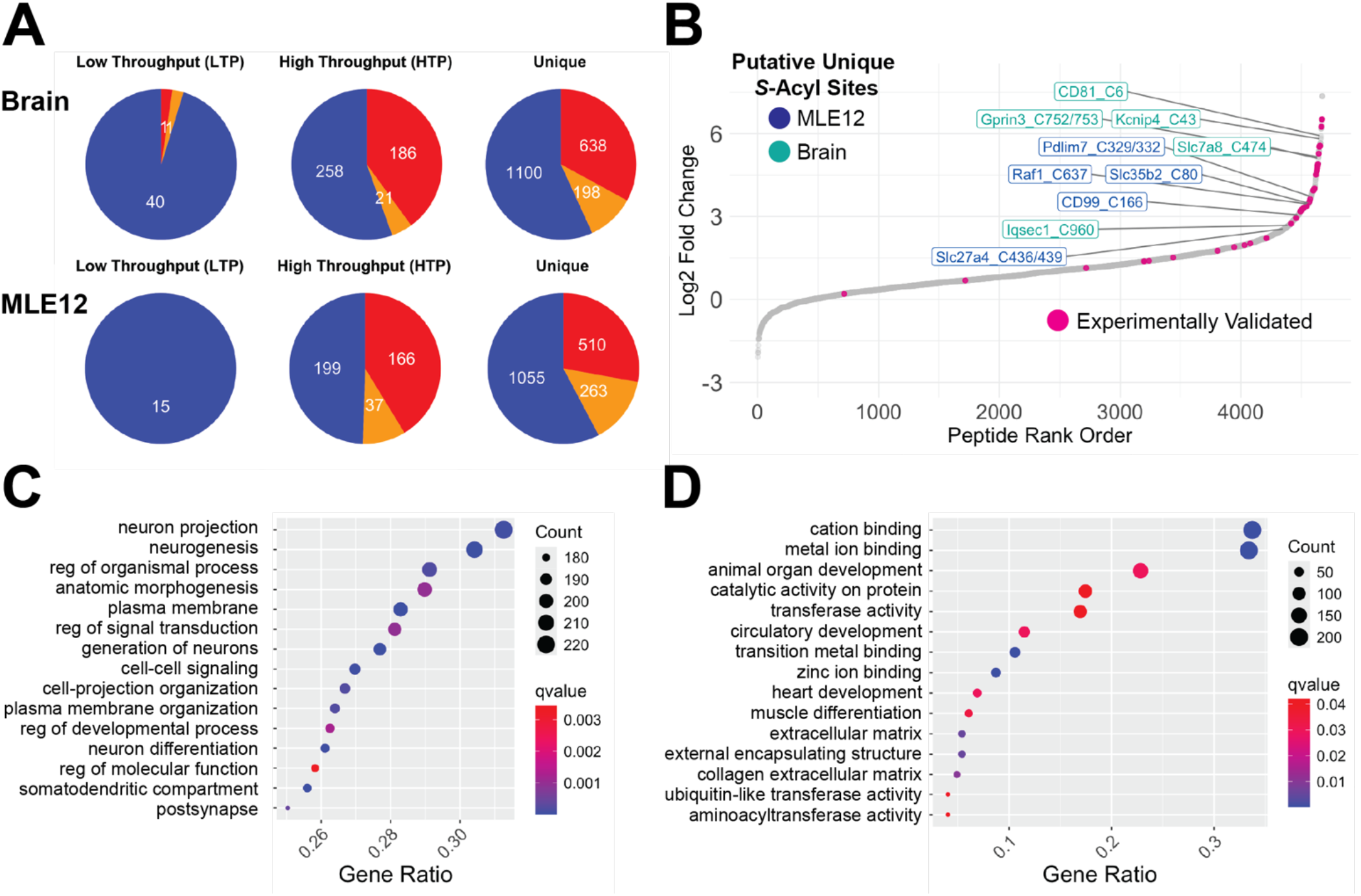
Palmitoyl database matching and GO analyses of Acyl-Trap results. (A) Pie charts of Brain and MLE-12 database site matches grouped by type (experimentally validated / low throughput (LTP), Swisspalm / high throughput (HTP) and unique) and colored based on Acyl-Trap experimental assignment: enriched high probability (blue, Log2FC ± NH_2_OH > 0.59 and adjusted p-value < 0.05, enriched low probability (orange, Log2FC ± NH_2_OH > 0.59 and adjusted p-value > 0.05) and unenriched (red, Log2FC ± NH_2_OH < 0.59) (L. (B) Rank order plot of all TMT-quantitated peptides from combined brain and MLE12 data sorted for Log2FC ± NH_2_OH. Highlighted in magenta are the peptides bearing *S*-palmitoyl sites matching to LTP data. Labels in blue and teal represent 5 selected novel putative *S*-acyl sites for MLE-12 and brain, respectively. (C and D) Over-representation analyses (package ClusterProfiler) comparing a list of enriched proteins to an experiment-derived universal gene list for (C) mouse brain and (D) MLE-12 cells.

Next, results from Acyl-Trap were matched to HTP data obtained from the entirety of Swisspalm (4381 mouse sites). In brain, 465 Cys sites matched to the HTP database with 279 sites demonstrating enrichment (258 high- and 21 low-probability, respectively) and 186 sites being assigned to unenriched. In MLE-12 cells, 402 Cys sites matched to the HTP database with 236 sites demonstrating enrichment (199 high- and 37 low-probability, respectively) and 166 sites being assigned to unenriched. There are a multitude of explanations for the lower degree of matching with HTP compared to LTP datasets, including the diverse array of experimental methods and assignment criteria included in HTP datasets. For example, while NH_2_OH-dependence is utilized in the vast majority of small-scale experiments and a criterion upon which Acyl-Trap performs *S*-acyl assignments, it is not a universal criterion for HTP assignment of *S*-acyl sites reported in Swisspalm.

### Allowing *N*-terminal Myristoylation as a Variable Modification Facilitates Identification of Dual Lipid Modified Sites

Protein *N*-terminal Gly residues may undergo *N*-terminal myristoylation (C12) facilitating transient membrane association and promoting *S*-palmitoylation of adjacent Cys residues^53^. Examples include certain Galpha isoforms (e.g., Gαi, Gαo)^54^ and Src family kinases^55^. The inclusion of protein *N*-term Gly *N*-myristoylation (+210.19836 Da) during database searching of mouse brain Acyl-Trap data identified known *S*-acylated sites Cys^3^ from Gαi1 and Cys^3^ from Gαi2 Cys^3^ with Log2FC (± NH_2_OH) of 4.90 and 1.89, respectively. In contrast, *N*-myristoyl peptides were not identified in MLE12 cells. Taken together, the inclusion of a variable *N*-terminal Gly *N*-myristoyl modification provides a very modest gain in identification of *S*-acyl sites.

### Identification of Putative *S*-acyl Sites in Brain and MLE12 cells

As shown in Figure 4A, the majority of sites demonstrating high probability enrichment are “unique” in that they are not represented in either LTP or HTP data from SwissPalm/UniProt (1100 in Brain, 1055 in MLE12). Given Acyl-Trap’s ability to identify well-established sites and its methodological distinction of using peptide-level (rather than protein-level) *S*-acyl enrichment, these unique putative *S*-acylated sites may represent newly discovered sites of endogenous *S*-acylation. Shown in Figure 4B is a rank order plot of combined data (MLE12 and Brain) overlaying matches to the LTP dataset (in magenta). Highlighted are peptides bearing previously unreported sites of *S*-acylation (5 each from brain and MLE12). Within brain, these include Cys960 of Iqsec1 (BRAG2), a guanine nucleotide exchange factor (GEF) for ARF family GTPases, which plays an important role in vesicle trafficking, synapse formation and neurodevelopment^56^. The presence of liposomes has shown to promote Iqsec GEF activity by up to 2000-fold^57^ suggesting that *S*-acylation could provide another level of regulation for Iqsec1 by either promoting membrane localization, GEF activity or both. Other notable putative *S*-acyl identifications include Cys^43^ on potassium channel Kcnip, Cys^6^ on tetraspanin CD81, Cys^474^ on amino acid transporter Slc7a8 and Cys^752/753^ on GPCR modulator Gprin3. In MLE-12 cells, notable enriched sites include Cys^637^ on proto-oncogene Raf1, Cys^166^ on CD99, Cys^436/439^ on fatty acid transporter Slc27a5, Cys^80^ on adenylyl sulfate transporter Slc35b2 and Cys^329/332^ on kinase-scaffolding protein Pdlim7. Collectively these findings demonstrate the utility of Acyl-Trap for probing *S*-acylated proteomes with relatively small quantities of biological material, identifying both established and previously unreported putative sites of *S*-acylation.

### Gene Ontology Analysis of Protein *S*-Acylation

To examine the ability of Acyl-Trap to probe functional roles of *S*-acylation in vivo, results from brain and MLE-12 cells were subjected to over-representation analysis (ORA) with clusterProfiler^43^. A list of enriched genes was compared to a reference “universal” gene list consisting of the enriched, unenriched and undetected (i.e. present in unbound but not observed in the bound) proteins. As shown in Figure 4C/4D, ORA from brain and MLE-12 suggest that *S*-acylation is involved in a wide array of pathways with an emphasis on cell differentiation, protein modifications and signal transduction. In brain, the identified terms match closely with established functions of *S*-acylation in the nervous system, including “neuron projection”, “neurogenesis” and “plasma membrane region”. The ORA analysis demonstrates that Acyl-Trap provides sufficient data to support GO analyses and that future applications of Acyl-Trap on unique biological samples may facilitate novel discovery of *S*-acylation’s *in vivo* roles.

### Conditions for Long-Term Storage of Neutral Hydroxylamine

The bulk of *S*-acyl assays utilizing neutral NH_2_OH recommend preparing fresh solution and bringing to neutral pH immediately prior to each use. Given the time-consuming nature of this step, we assessed the feasibility of using frozen NH_2_OH for thioester cleavage. In the absence of EDTA, NH_2_OH has been shown to exhibit drastically diminished efficacy for *S*-acyl (i.e. thioester) cleavage^58^, suggesting the importance of transition metal chelation in stabilizing NH_2_OH. Neutral 2M NH_2_OH was therefore prepared in the presence of 5 mM EDTA and stored at -80 °C for up to 6 months. Aliquots were subjected to colorimetric titration with 4-nitrophenylacetate (4-NPA) which is known to undergo nucleophilic reaction with NH_2_OH to release 4-nitrophenolate^59^ (Figure S8A). Stored vs fresh solutions of NH_2_OH demonstrated identical reactivity to a 5-fold molar excess of 4-NPA (Figure S8B) and identical results when assessing H-Ras *S*-acylation by Acyl-RAC (Figure S8C). Thus, 2M NH_2_OH can be stored at -80 °C for at least 6 months in the presence of 5 mM EDTA without any appreciable loss in assay efficacy. Given the time-consuming nature of fresh hydroxylamine preparation and pH adjustment to neutral range, the described storage conditions should facilitate all *S*-acyl assays that utilize NH_2_OH.

## CONCLUSIONS

Since its introduction by Zougman et al in 2014^33^, suspension trapping has become widely adopted for bottom-up proteomic sample processing. The method is simple, amenable to small sample quantities and allows for simultaneous processing of many samples. Here we provide a simple and cost-effective synthetic protocol for thiol-reactive quartz (PDQ) and apply it for *S*-acyl enrichment of well-established *S*-acylated proteins and discovery of novel putative *S*-acyl sites from microgram quantities of input material. We show that suspension traps can support intact protein manipulation, including “on trap” labeling and elution, uncovering the potential for these devices to be adapted to basic laboratory procedures and “top-down” mass spectrometry workflows. It is also notable that the unbound protein or peptide fractions can be recovered, as this simplifies the concomitant analysis of the global (non-targeted, non-enriched) proteome and S-acylome of the same sample.

Chemoproteomics is increasingly performed in large scale, aided by techniques such as single-pot, solid-phase-enhanced sample-preparation (SP3)^60^, clickable probes^61^ and multi/hyperplexing^62,63^. Similarly, compared to ABE and Acyl-RAC, Acyl-Trap helps overcome the “throughput barriers” associated with traditional protein precipitation and recovery. While suspension trapping has been used in series with phosphopeptide enrichment^64^, and for on-trap isotopic Cys labeling^65^, our demonstration of its compatibility with thioester cleavage and in situ Cys capture, at the protein and peptide level, serves as a proof-of-principle for the utility of suspension trapping in covalent PTM-enrichment. Given the wide range of commercially available organosilanes, a future toolbox of “chemically tailored” suspension traps may facilitate a range of targeted enrichments and allow PTMs (including *S*-acylation) to become a component of standardized (chemo)proteomic workflows for drug discovery and large scale ‘omics.

Finally, there are limitations of Acyl-Trap that should be noted. Similar to ABE and Acyl-RAC, Acyl-Trap does not reveal molecular details of the *S*-acyl moiety itself. While *S*-palmitoylation is a highly studied and perhaps most prevalent type of *S*-acylation, inferring the exact type of *S*-acyl moiety should be performed within a biological context and may benefit from complementary approaches such as bioorthogonal lipid labeling^25-28^. These assays may also not be well-suited for differentiating a single vs multiple *S*-acyl groups per protein, which has been elegantly overcome with PEG-based mass shifting for the study of individual poly-*S*-acylated proteins^58^. In addition, the studies herein used less than one-fourth of the 18 available TMT channels for the +NH_2_OH samples thus may not be fully realizing the potential of Acyl-Trap for maximal coverage of the *S*-acylome. Similar to other *S*-Acyl assays, our coverage is limited to those peptides identified by trypsinolysis, which may be overcome by the use of orthogonal proteases^66,67^.

## DATA AVAILABILITY STATEMENT

Raw and processed data, and metadata, are available at the ProteomeXchange Consortium (PXD050341) via the MassIVE repository (MSV000094234; ftp://massive.ucsd.edu/v07/MSV000094234/). All related code is available at https://github.com/mt-forrester/acyl_trap_jpr. Adapters needed to perform Acyl-Trap with microfuge tubes can be 3D-printed with .stl file: https://www.thingiverse.com/thing:6423695.

## SUPPORTING INFORMATION

Supplementary Methods; Figure S1. Synthesis and overview of PDQ traps; Figure S2. Detailed assembly of PDQ Traps; Figure S3. Evaluation of experimental conditions unique to Acyl-Trap; Figure S4. Cysteine enrichment efficiency; Figure S5. Under- and over-labeling with on-quartz TMT labeling; Figure S6. Coefficient of variations from Acyl-Trap experiments; Figure S7. Prevalence of methionine oxidation; Figure S8. Conditions for long-term hydroxylamine storage; Figure S9. Entire images of blots and SDS-PAGE gels; Table S1. Isobaric quant data from brain and MLE12 Acyl-Trap experiments (.xlsx).

## ACKNOWLEDGEMENTS

We thank Erik J. Soderblom (Duke Proteomics and Metabolomics Core Facility), Christine E. Eyler (Department of Radiation Oncology), Patrick J. Casey (Duke Pharmacology and Cancer Biology) and John R. Wilson (Protifi, LLC) for helpful advice and discussions. The work was supported in part by an award from the Duke University School of Medicine and NIH R41HD105560 (to M.W.F), the Duke Office of Physician Scientist Development (OPSD) and T32HL160494 (to M.T.F.) and research awards from NHLBI/NIH (R01HL160939, and R01HL153375) to P.R.T.

## CONFLICT OF INTEREST STATEMENT

M.T.F., J.R.E, P.R.T, and A.T. have no conflicts of interest. M.W.F. is the principal investigator on a sponsored research agreement between Duke University and Circumvent Pharmaceuticals, Inc.

## Notes

https://proteomecentral.proteomexchange.org/cgi/GetDataset?ID=PXD050341

ftp://massive.ucsd.edu/v07/MSV000094234

https://github.com/mt-forrester/acyl_trap_jpr

https://www.thingiverse.com/thing:6423695

### Summary of Updates

Addition of H-Ras Cys181Ser/Cys184Ser mutant data, entire blots per journal requirements, typographical corrections, change of "S-palmitoylation" to "S-acylation" when indeterminate acyl group.

