## Supporting Information for "Analysis of Protein Cysteine Acylation Using a Modified Suspension Trap (Acyl-Trap)"

Supplementary Methods 2

Figure S1. Synthesis and overview of PDQ traps 4

Figure S2. Detailed assembly of PDQ Traps 5

Figure S3. Evaluation of experimental conditions unique to Acyl-Trap 6

Figure S4. Cysteine enrichment efficiency 7

Figure S5. Under- and over-labeling with on-quartz TMT labeling 8

Figure S6. Coefficient of variations from Acyl-Trap experiments 9

Figure S7. Prevalence of methionine oxidation 10

Figure S8. Conditions for long-term hydroxylamine storage 11

Figure S9. Entire images of blots and SDS-PAGE gels 12

Table S1. Isobaric quant data from brain and MLE12 Acyl-Trap experiments (.xlsx)

SUPPLEMENTARY METHODS

*3D Printing of trap-to-microfuge adapters* – Performing Acyl-Trap with pipette tip traps (as herein described) requires adapting pipette tips into microfuge tubes to support centrifugation. We have designed and printed these adapters with poly-lactic acid on an Ultimaker S3 printer. Further information and related .stl files are available through Thingiverse by searching for “Pipette Tip - Microfuge Adapter”, 6423695, or via the following link: https://www.thingiverse.com/thing:6423695.

*Thiol assay by DTNB reaction (Ellman’s reagent)* – Mk360 filters were subjected to silanization with (3-mercaptopropyl)trimethoxysilane as in the Main Text and repeatedly washed with xylenes and isopropanol. Thiol-derivatized filters were cut into quarters and mixed with 50 mM HEPES, 2 mM EDTA, pH 7.5 containing 0.5 mM DTNB (from a 25 mM stock solution in EtOH). Supernatant was collected and absorbance measured at 405 nm. Standards were cysteine-HCl salt freshly prepared in 50 mM HEPES, 2 mM EDTA, pH 7.5.

*Trapping efficiency of Mk360 quartz at neutral pH* – Cultured MLE-12 cells were lysed by probe sonication in 50 mM HEPES, 2 mM EDTA containing either 2.5% SDS or 6 M Gdn-Cl followed by centrifugation at 2,000 *xg* for 5 min and measurement of protein concentration by detergent-compatible Bradford assay. Twenty micrograms of lysate in 50 µL of either 2.5% SDS or 6 M Gdn-Cl were precipitated with the indicated volume of EtOH or MeOH for 10 min at room temp. Samples were loaded to a quartz trap (non-PDQ), centrifuged at 3000 *xg*, then washed four times with MeOH. To the trapped protein was added 100 µL of Pierce Dilution-Free Gold BCA reagent. After 5 min at room temp, traps were centrifuged at 3000 *xg* for 1 min and eluant was collected. An additional 100 µL of Pierce Dilution-Free Gold BCA reagent was added and again collected after 5 min at room temp. Eluants were subjected to absorbance measurement at 492 nm along with BSA standards. Note that the BCA reaction was used to measure trapped protein as it does not exhibit any color change when exposed to quartz, whereas Bradford reagents demonstrated color change. Non-derivatized quartz was used in lieu of PDQ to ensure no potential interference of the pyridyl disulfide groups on the BCA assay.

*Colorimetric assay of trypsin activity in the presence of $NH_2OH$* – Reactions (1 ml each) of sequencing grade trypsin, BAPNA and ratios of NaCl and $NH_2OH$ (to maintain equal osmolarity in each reaction) in 100 mM HEPES, 5 mM EDTA, pH 7.2 were allowed to react at 37 °C. Reactions contained 2.5 µL of trypsin (from 1 mg/mL stock) and 2 µl of BAPNA (from a 200 mM stock in DMSO). Aliquots of 150 µL were removed, transferred to a 96 well plate and absorbance measured at 405 nm at indicated timepoints.

*Determination of Cys-enrichment efficiency* – Fragpipe output ion files (containing precursor ions) were read into RStudio and stringr package used to identify the presence of Cys-containing precursors via searching for “C” in column ‘Peptide Sequence’.

*Evaluation of Met oxidation and TMT under/over-labeling* – Data were searched in Fragpipe to allow for identification of precursors lacking a TMT modification of peptide N-terminus or Lys sidechain (under-labeling) or TMT modification on Ser, Thr, Tyr or His side chains (over-labeling) in a manner similar to Zecha et al[1]. For under-labeling, variable mods were N-term TMT (+304.20715 Da), Lys TMT (+304.20715 Da), Met oxidation (+15.9949 Da), Cys NEM adduct (+125.047676 Da) and Cys carboxymethylation (+57.02146 Da). For over-labeling, N-term and Lys TMT mods were set to fixed and TMT mod (+304.20715 Da) was allowed as variable on Ser, Thr, His or Tyr. Fragpipe ion outputs were read into RStudio and TMT labeling positions were extracted using StringR package searching for presence or absence of Lys residues, as well as the TMT mass tag.

*Acyl-RAC Assay of HEK293 cell Lysate and Transfected H-Ras* – A 6 cm dish of HEK293 cells was transfected with 4 μg of pCDNA3.1-3xHA-H-Ras using 3:1 ratio (μL/μg) of PEI. After 24h, cells are collected with PBS and subjected to Acyl-RAC as described[2] with several modifications as outlined below. The HEK293 cell pellet was lysed in 800 μL of 50 mM HEPES, 2 mM EDTA, 0.5% Triton X-100 containing 20 mM NEM via probe sonication. Crude lysate was centrifuged at 2000 *xg* for 10 min. Supernatant was diluted to 2 mL with 100 mM HEPES, 2 mM EDTA and SDS was added to final 1% (w/v) concentration. Additional NEM was added to maintain 20 mM concentration during blocking. Blocking was performed at 50 °C for 30 min. Three volumes of room temp MeOH was added, agitated and placed to -20 °C for 30 min. Precipitated protein was collected by centrifugation at 3000 *xg* for 5 min. Supernatant was removed via suction through 21G needle and pellet was vortexed in 6 ml of MeOH to ensure efficient removal of NEM without loss of protein. This was again centrifuged at 3000 *xg* for 5 min, pellet resuspended in 1 mL of 100 mM HEPES, 5 mM EDTA, 1% SDS and split into two equal fractions (450 μL each) onto 30 μL of pyridyl disulfide Sepharose (PDS). To one fraction was added $NH_2OH$ (from frozen 4M stock) to final concentration of 500 mM and to the other was added water. Binding reactions were rotated at room temp overnight (16 h), washed with 1 mL of 50 mM HEPES, 2 mM EDTA, 1% SDS four times, eluted with 50 μL of was buffer containing 20 mM DTT at room temp for 15 min, then subjected to SDS-PAGE and Western Blotting.

*Spectrophotometric evaluation of 4-NPA cleavage by $NH_2OH$* – Aliquots of frozen or freshly prepared $NH_2OH$ (2 or 4M) in 5 mM EDTA were diluted into 100 mM phosphate buffer pH 6.0 containing 5 mM EDTA to a final concentration of 2 mM $NH_2OH$, followed by immediate addition of 4-NPA (prepared as a 1M stock in DMSO) to final concentration of 10 mM 4-NPA. To control reactions was added either water in lieu of $NH_2OH$ or DMSO in lieu of 4-NPA. Reactions (1 mL each) were allowed to proceed at room temp for 30 min, then 300 μL was transferred to a 96 well plate to measure absorbance at 405 nm.

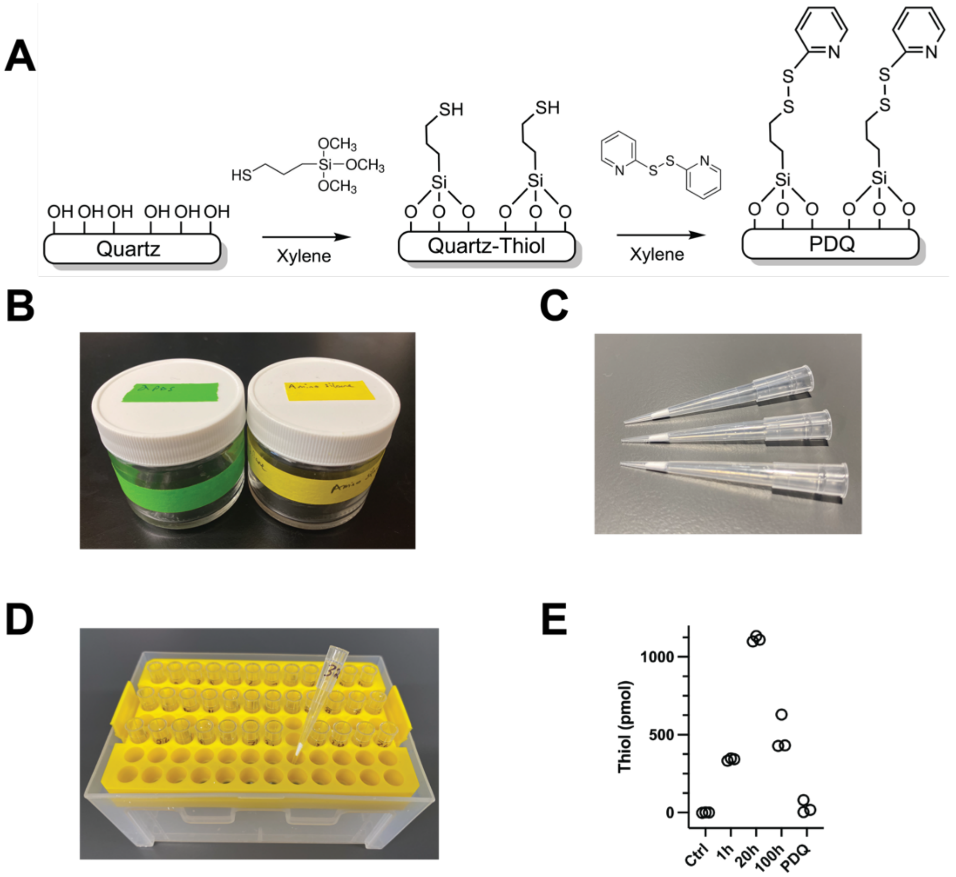


**Figure S1**. Synthesis of pyridyl disulfide quartz (PDQ) and creation of PDQ traps. (A) Chemical derivatization of quartz to PDQ. The two-step process involves "silanization" of quartz with (3-mercaptopropyl)trimethoxysilane followed by thiol oxidation to the 2-pyridyl disufide quartz (PDQ). Two adjacent derivatizations are illustrated. (B) Separate TraceClean® glass containers used for silanization and thiol oxidation reactions. (C) Immobilized PDQ traps ready for use. (D) PDQ traps in a 96-well centrifugation apparatus facilitating multiplexing and simultaneous processing of multiple samples. (E) Thiol measurement by DTNB reaction of Mk360 quartz during PDQ synthesis process. Shown on the x-axis are timepoints for the silanization reaction compared to unreacted (control) or PDQ (which has undergone thiol oxidation with 2-pyridyl disulfide). For each measurement, n =3. Measurements were at 405 nm with freshly prepared Cysteine-HCl as thiol standards.

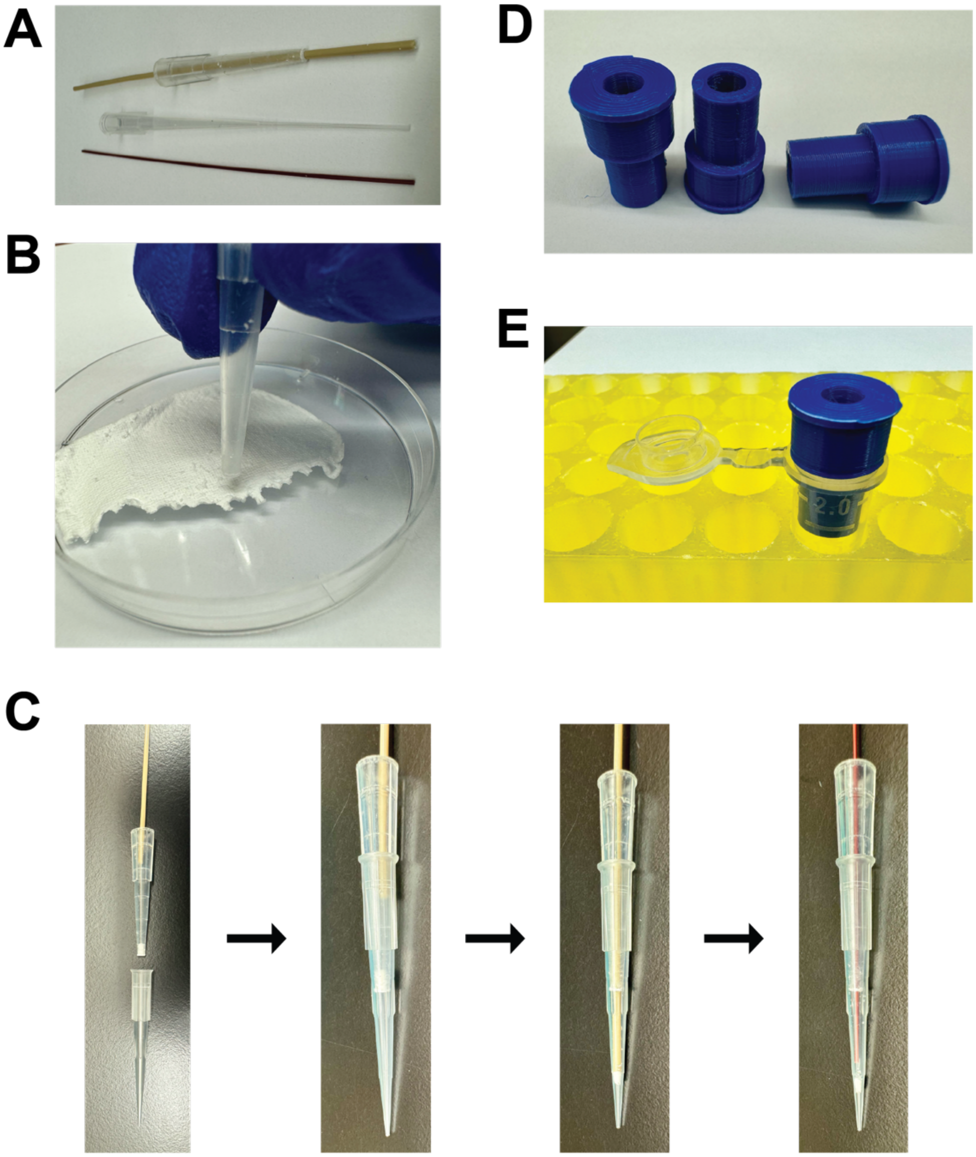


Figure S2. Detailed assembly of PDQ traps and single-trap centrifugation apparatus. (A) Basic tools include a blunt P200 tip, rigid capillary tubing and a loading tip for SDS-PAGE (optional in lieu of thin capillary tubing). (B) Blunt tip used to acquire "cores" of PDQ via downward pressure and twisting. (C) Loading of the P200 tip first with thick capillary tubing then fully seated with thin tubing. This process is repeated 3 times. (D) Centrifugation adapters 3-D printed in poly-lactic acid that house the P200 PDQ trap in microfuge tubes. (E) Adapter housed in a 2.0 ml microfuge tube. The PDQ trap seats into the apparatus.

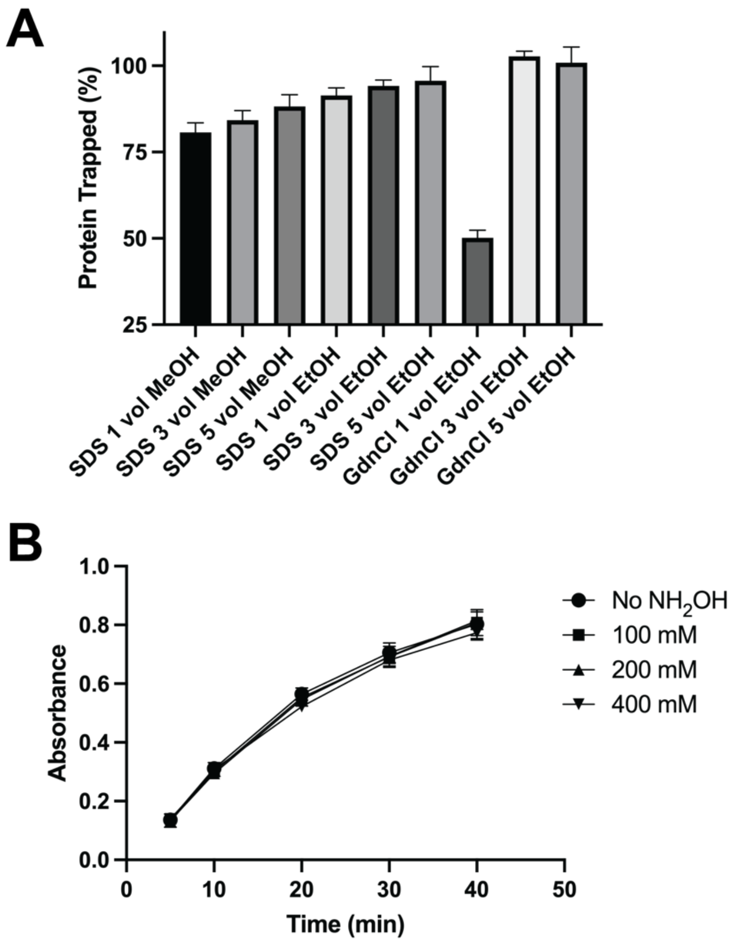


**Figure S3**. Evaluation of experimental conditions unique to Acyl-Trap. (A) Protein trapping efficiency at neutral pH. Lysates of MLE-12 cells (20 µg each) were subjected to precipitation with the indicated volume of EtOH or MeOH for 10 min at room temp, trapped as per Acyl-Trap protocol and protein measured with Pierce BCA Gold reagent as described in the methods. For each condition, n = 4. (B) Trypsin activity in the presence of $NH_2OH$. Trypsin (2.5 ng/mL) was incubated with colorimetric trypsin substrate BAPNA (0.4 mM) at 37 °C for indicated timepoints and absorbance measured at 405 nm. For each condition, n = 4.

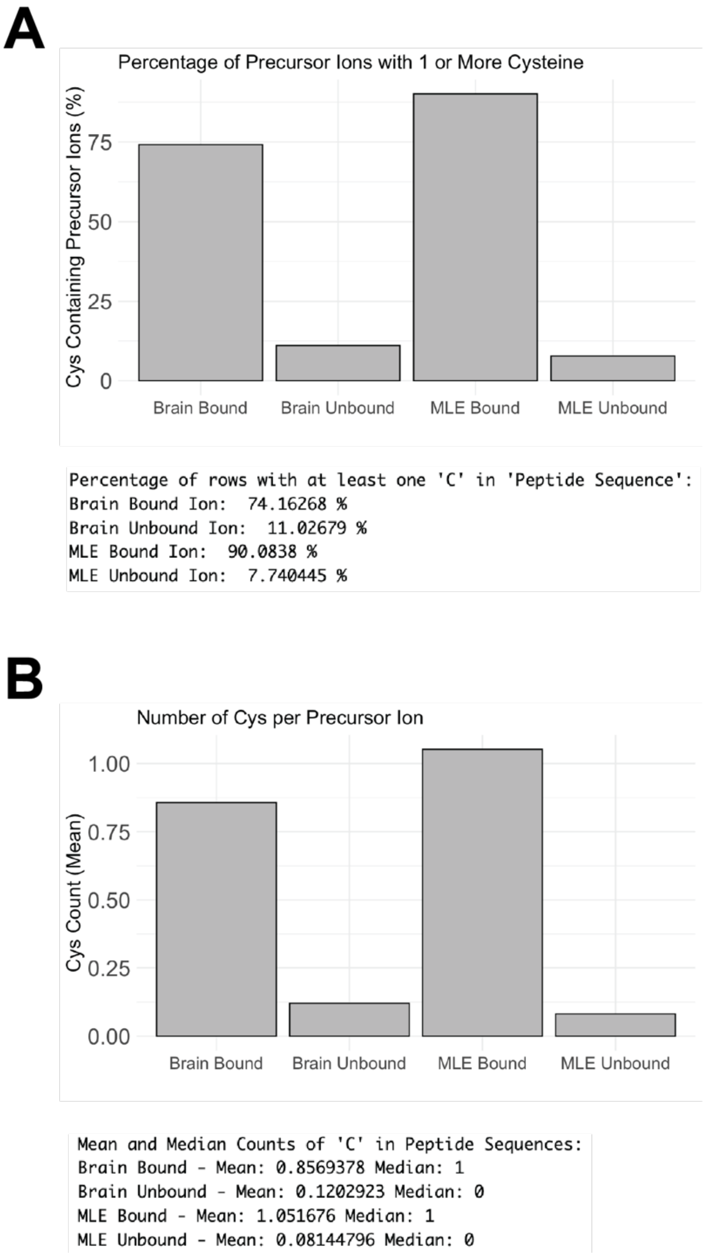


**Figure S4**. Cysteine enrichment efficiency in Acyl-Trap. (A) Percentage of precursor ions containing at least one Cys residue. (B) Mean number of Cys per precursor ion. Comparisons are between bound and unbound for each sample type (brain or MLE12 cells).

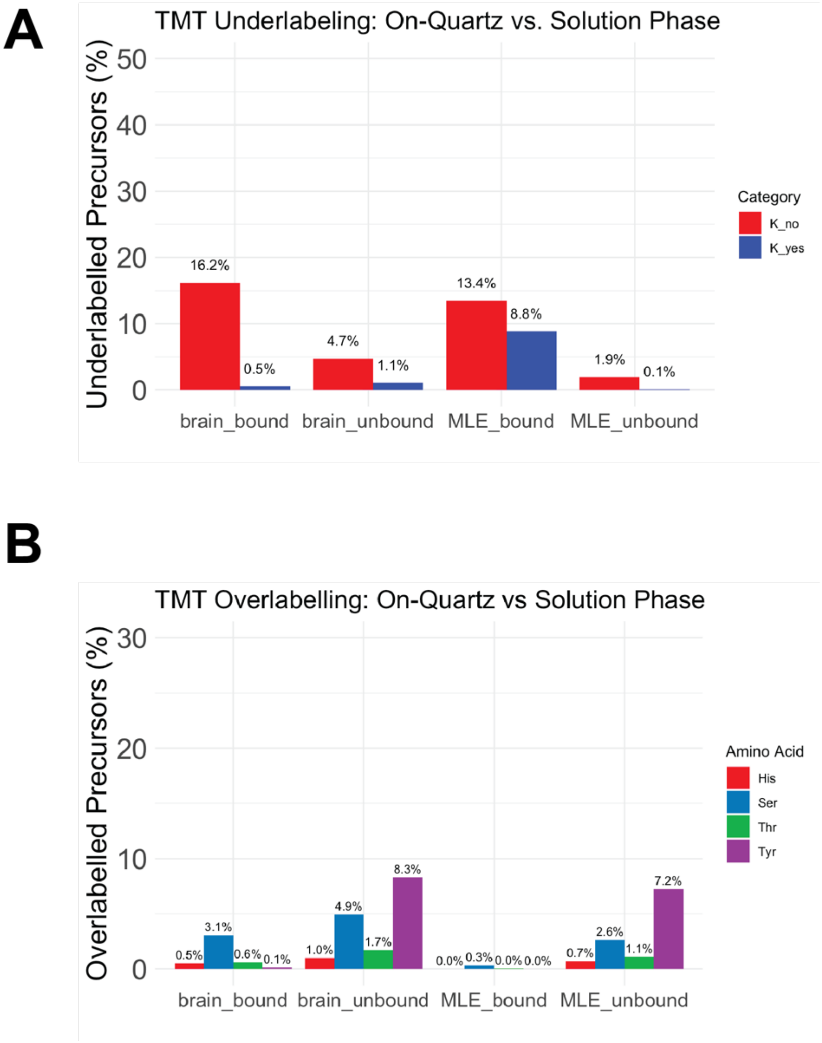


**Figure S5**. Under- and over-labeling with on-quartz TMT labeling in Acyl-Trap. (A) Percentage of precursor ions lacking TMT modification on Lys-containing or non-Lys-containing (i.e., Arg-containing) peptides. (B) Percentage of precursor ions with fixed Lys and peptide N-terminal TMT modification that also possess TMT modification on His, Ser, Thr or Tyr side chains. Bound samples were TMT labeled on-quartz (while covalently immobilized to PDQ). Unbound samples were labeled while bound to C18 stage tip as described by Myers et al.[3]

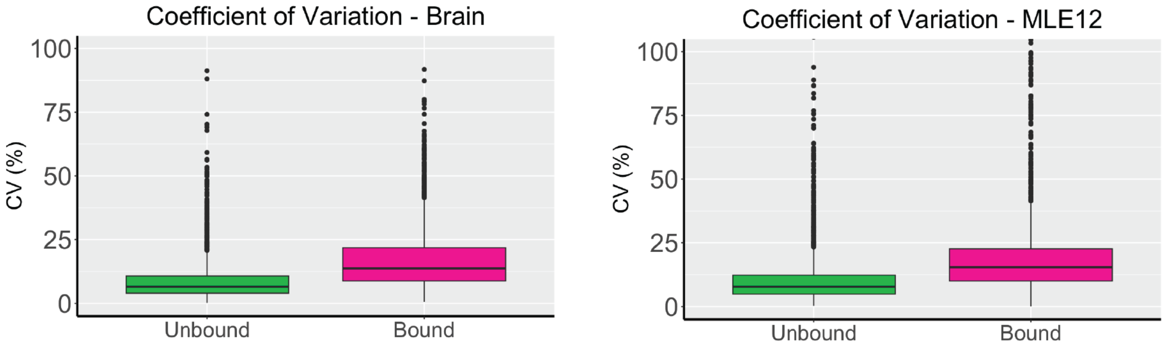


| type <fctr> | brain_median_CV <dbl> |
|---|---|
| Unbound | 6.55541 |
| Bound | 13.70243 |

| type <fctr> | MLE_median_CV <dbl> |
|---|---|
| Unbound | 7.809699 |
| Bound | 15.420780 |

**Figure S6**. Coefficient of variations from Acyl-Trap experiments. Post-processed data were subjected to reverse log2 transformation and calculation of CV across the minus and plus $NH_2OH$ groups.

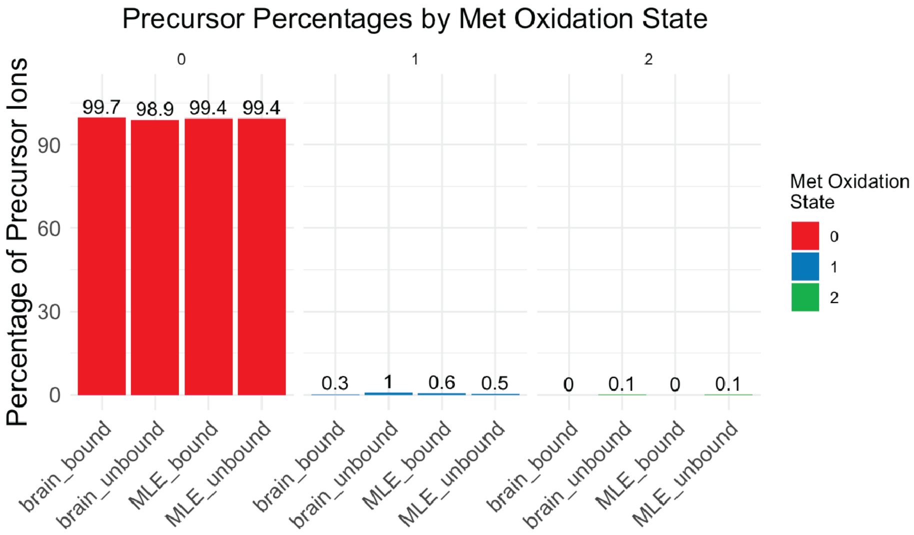


**Figure S7**. Precursor ions containing oxidized Met. MSFragger search was performed with Met oxidation as a variable modification (max 4 per peptide). Precursor data from sample types (indicated on x axis) were then analyzed for the presence of Met oxidations and shown as a percentage of all precursor ions within that sample.

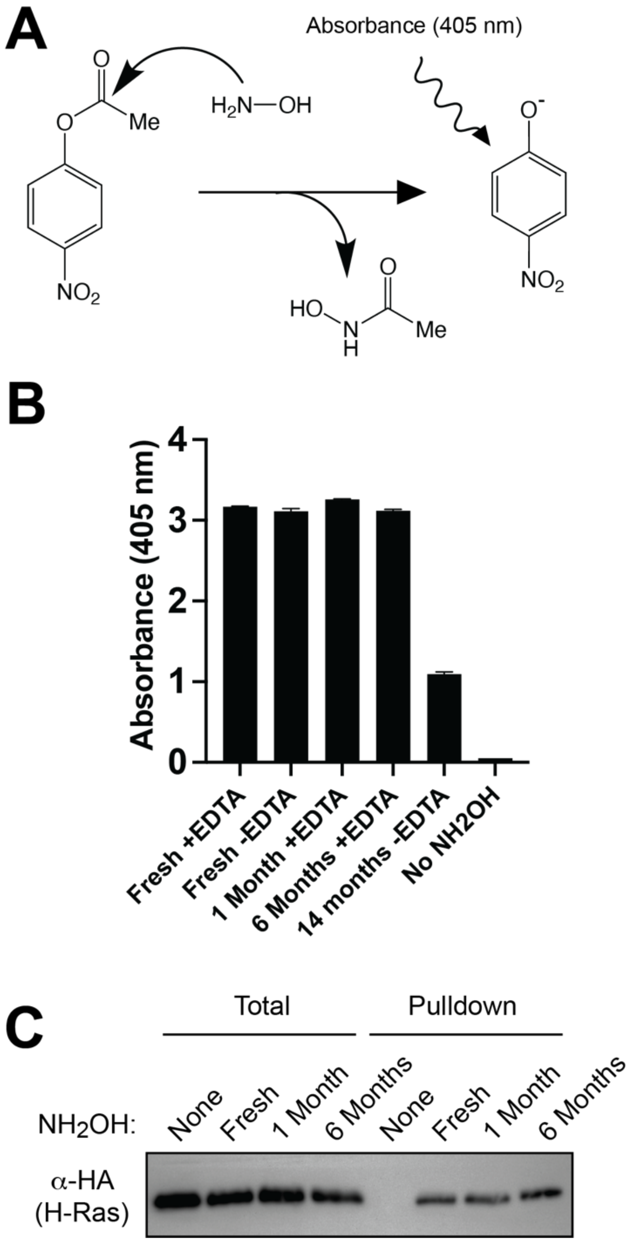


**Figure S8**. Conditions for long-term hydroxylamine storage compatible with *S*-acyl assays. (A) Nucleophilic cleavage of 4-nitrophenyl acetate (4-NPA) by $NH_2OH$ leads to release of 4-nitrophenolate[4], which exhibits a strong absorbance at 405 nm. (B) Hydroxylamine stock solutions (2 M) were diluted to 2 mM in 100 mM phosphate with 5 mM EDTA (pH 6.0) followed by addition of 10 mM 4-NPA and incubation at RT for 30 min. (C) HEK293 cells transfected with HA-tagged H-Ras were subjected to Acyl-RAC employing either fresh or stored $NH_2OH$. Western blot is with anti-HA antibody. Data shown are representative from n=4 experiments.

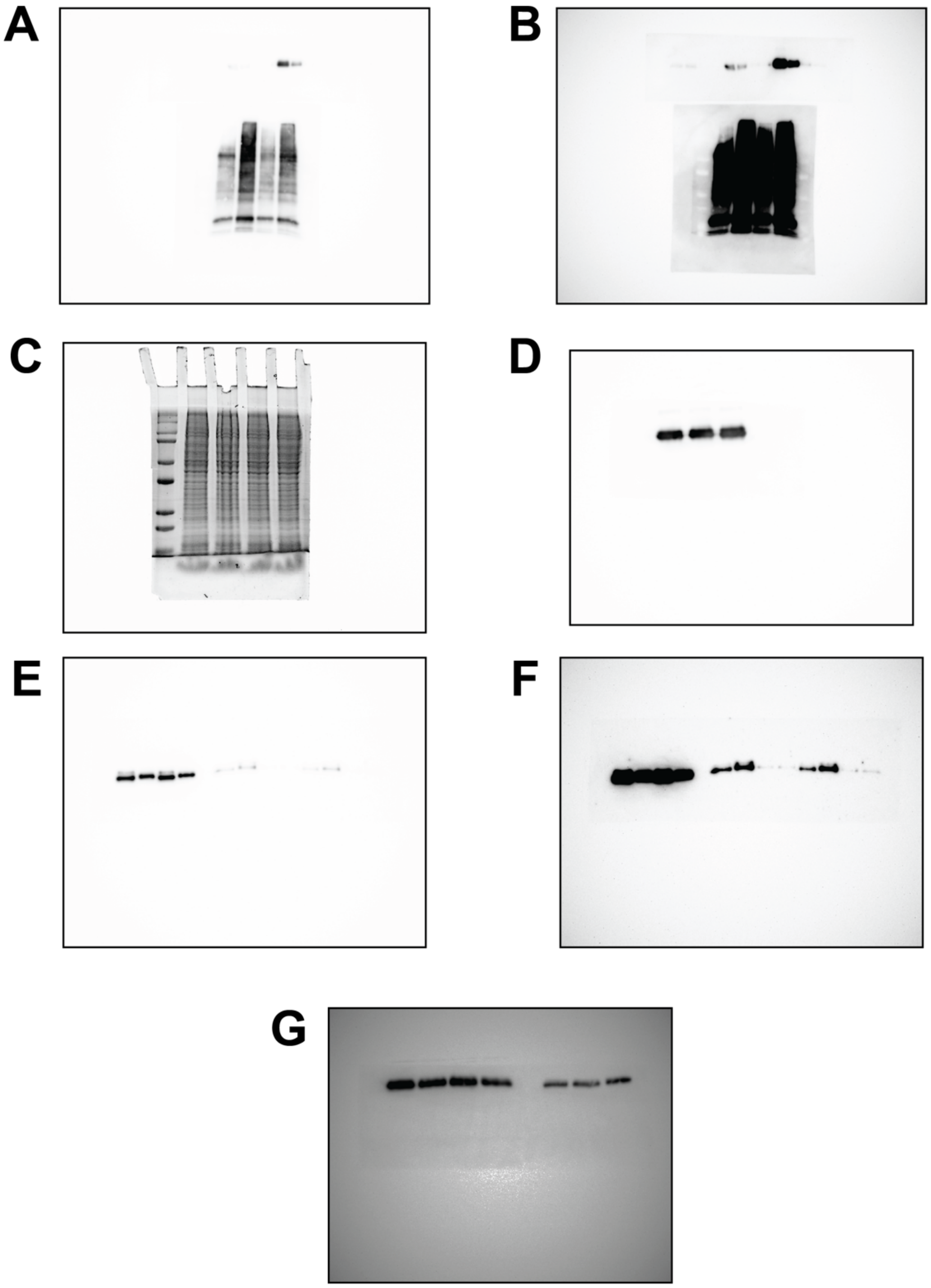


**Figure S9**. Entire images of western blots and associated data (e.g., SDS-PAGE + Coomassie). (A) Avidin HRP and α-HA blot used in Figure 2A and 2C, respectively. (B) Same as (A) but different exposure. (C) Coomassie stain used in Figure 2A. (D) α-HA loading control (input) used in Figure 2C. (E) α-HA blot used in Figure 2D. (F) Same as (E) but different exposure. (G) α-HA blot from Figure S8C showing $NH_2OH$ efficacy with storage.

SUPPLEMENTARY TABLE

**Table S1**. Isobaric quant data from brain and MLE12 Acyl-Trap experiments. Bound and unbound datasets are separated by tabs. All replicate quant data are log2 transformed. For the bound data, each row represents an individual Cys site indicated in the column "cys_experiment". Low throughput (LTP) matches include all rows that have yes in either column "uniprot_match" or "swisspalm_validated_match". High throughput (HTP) matches include all rows with "yes" in column "swisspalm_all_match". These results are summarized in column "database_site_match". Unbound data are arranged similarly except each row represents a protein instead of a Cys site and columns for database site matches are not applicable.
